# Slo2 potassium channel function depends on a SCYL1 protein

**DOI:** 10.1101/864645

**Authors:** Long-Gang Niu, Ping Liu, Zhao-Wen Wang, Bojun Chen

## Abstract

Slo2 potassium channels play important roles in neuronal function, and their mutations in humans cause epilepsies and cognitive defects. However, little is known how Slo2 function is regulated by other proteins. Here we found that the function of *C. elegans* Slo2 (SLO-2) depends on *adr-1*, a gene important to RNA editing. However, *slo-2* transcripts have no detectable RNA editing events and exhibit similar expression levels in wild type and *adr-1* mutants. In contrast, mRNA level of *scyl-1*, which encodes an orthologue of mammalian SCYL1, is greatly reduced in *adr-1* mutants due to deficient RNA editing at a single adenosine in its 3’-UTR. SCYL-1 physically interacts with SLO-2 in neurons. Single-channel open probability of SLO-2 in neurons is reduced by ∼50% in *scyl-1* knockout whereas that of human Slo2.2/Slack is doubled by SCYL1 in a heterologous expression system. These results suggest that SCYL-1/SCYL1 is an evolutionarily conserved regulator of Slo2 channels.

## Introduction

Slo2 channels are large-conductance potassium channels existing in mammals as well as invertebrates (Kaczmarek, 2013; Yuan et al., 2000). They are the primary conductor of delayed outward currents in many neurons examined (Budelli et al., 2009; Liu et al., 2014). Human and mouse each has two Slo2 channels (Slo2.1/Slick and Slo2.2/Slack) (Kaczmarek, 2013), whereas the nematode *C. elegans* has only one (SLO-2). These channels are abundantly expressed in the nervous system (Bhattacharjee et al., 2002; Bhattacharjee et al., 2005; Joiner et al., 1998; Liu et al., 2018; Rizzi et al., 2016), and play major roles in shaping neuronal electrical properties and regulating neurotransmitter release (Kaczmarek, 2013; Liu et al., 2014). Mutations of Slo2 channels cause epilepsies and severe intellectual disabilities in humans (Ambrosino et al., 2018; Cataldi et al., 2019; Evely et al., 2017; Gururaj et al., 2017; Hansen et al., 2017; Kawasaki et al., 2017; Lim et al., 2016; McTague et al., 2018; Rizzo et al., 2016), and reduced tolerance to hypoxic environment in worms (Yuan et al., 2003). Emerging evidence suggests that physiological functions of these channels depend on other proteins. For example, in mice, the fragile mental retardation protein (FMRP), a RNA binding protein, enhances Slack activity by binding to its carboxyl terminus (Brown et al., 2010). In worms, HRPU-2, a RNA/DNA binding protein, controls the expression level of SLO-2 through a posttranscriptional effect (Liu et al., 2018).

RNA editing is an evolutionally conserved post-transcriptional process catalyzed by ADARs (*a*denosine *d*eaminases *a*cting on *R*NA) (Gott and Emeson, 2000; Jin et al., 2009). ADARs convert adenosine (A) to inosine (I) in double-stranded RNA. Since inosine is interpreted as guanosine (G) by cellular machineries (Basilio et al., 1962), A-to-I RNA editing may alter the function of a protein by changing its coding potential, or regulate gene expression through altering alternative splicing, microRNA processing, or RNA interference(Deffit and Hundley, 2016; Nishikura, 2016). Human and mouse each has three ADARs: ADAR1, ADAR2 and ADAR3 (Chen et al., 2000; Kim et al., 1994; Melcher et al., 1996). ADAR1 and ADAR2 possess deaminase activity and catalyzes the A-to-I conversion (Tan et al., 2017), whereas ADAR3 is catalytically inactive with regulatory roles in RNA editing (Nishikura, 2016) Millions of A-to-I editing sites have been detected in the human transcriptome through RNA-seq, with the vast majority of them found in non-coding regions (Nishikura, 2016). Biological effects of RNA editing at coding regions have been revealed for a variety of genes, including those encoding ligand- and voltage-gated ion channels and G protein-coupled receptors (Bhalla et al., 2004; Brusa et al., 1995; Burns et al., 1997; Gonzalez et al., 2011; Huang et al., 2012; Lomeli et al., 1994; Palladino et al., 2000; Rula et al., 2008; Sommer et al., 1991; Streit et al., 2011). However, little is known about the roles of RNA editing in non-coding regions (Nishikura, 2016).

In a genetic screen for suppressors of a sluggish phenotype caused by expressing a hyperactive SLO-2 in worms, we isolated mutants of several genes, including *adr-1*, which encodes one of two ADARs in *C. elegans* (ADR-1 and ADR-2). While ADR-2 has deaminase activity and plays an indispensable role in the A-to-I conversion, ADR-1 is catalytically inactive but can regulate RNA editing by binding to selected target mRNA and altering the accessibility of specific adenosines to ADR-2 (Ganem et al., 2019; Rajendren et al., 2018; Washburn et al., 2014). We found that loss-of-function (*lf*) mutations of *adr-1* impairs SLO-2 function through altering RNA editing of *scyl-1*,which encodes an orthologue of human and mouse SCYL1. In *adr-1(lf)* mutants, a lack of A-to-I conversion at a specific site in *scyl-1* 3’-UTR causes reduced *scyl-1* expression. Knockout of *scyl-1* severely reduces SLO-2 current in worms while coexpression of SCYL1 with human Slack in *Xenopus* oocytes greatly augments channel activity. These results suggest that SCYL-1/SCYL1 likely plays an evolutionarily conserved role in physiological functions of Slo2 channels. Mutations or knockout mammalian SCYL1 may cause neural degeneration, intellectual disabilities, and liver failure, but the underlying mechanisms are unclear (Lenz et al., 2018; Li et al., 2019; Shohet et al., 2019; Spagnoli et al., 2018, 2019). The revelation of SCYL-1/SCYL1 as a protein important to Slo2 channels suggests a potential link between diseases caused by SCLY1 mutations and Slo2 channel functions.

## Results

### adr-1 mutants suppress sluggish phenotype of slo-2(gf)

In a genetic screen for mutants that suppressed a sluggish phenotype caused by an engineered hyperactive or gain-of-function (*gf*) SLO-2 (Liu et al., 2018), we isolated two mutants (*zw80* and *zw81*) of the *adr-1* gene, as revealed by analyses of whole-genome sequencing data. *zw80* and *zw81* carry nonsense mutations leading to premature stops at tryptophan (W) 366 and W33, respectively (**Fig. 1A**). *slo-2(gf)* worms showed greatly decreased locomotion speed compared with wild type, and this phenotype was substantially alleviated in *slo-2(gf);adr-1(lf)* double mutants (**Fig. 1B**). To confirm that the suppression of *slo-2(gf)* phenotype resulted from mutations of *adr-1* rather than that of another gene, we created a new *adr-1* mutant allele (*zw96*) by introducing a premature stop codon at serine (S) 333 (**Fig. 1A**) using the CRISPR/Cas9 approach. The sluggish phenotype of *slo-2(gf)* was similarly suppressed by *adr-1(zw96)*, which, by itself, did not enhance locomotion speed (**Fig. 1B**). Expression of wild-type *adr-1* under the control of the pan-neuronal *rab-3* promotor (P*rab-3*) in *slo-2(gf);adr-1(zw96)* reinstated the sluggish phenotype (**Fig. 1B**). These results indicate that the sluggish phenotype of *slo-2(gf)* is mainly caused by SLO-2 hyperactivity in neurons, and that neuronal function of SLO-2(*gf*) depends on ADR-1.

**Figure 1.**
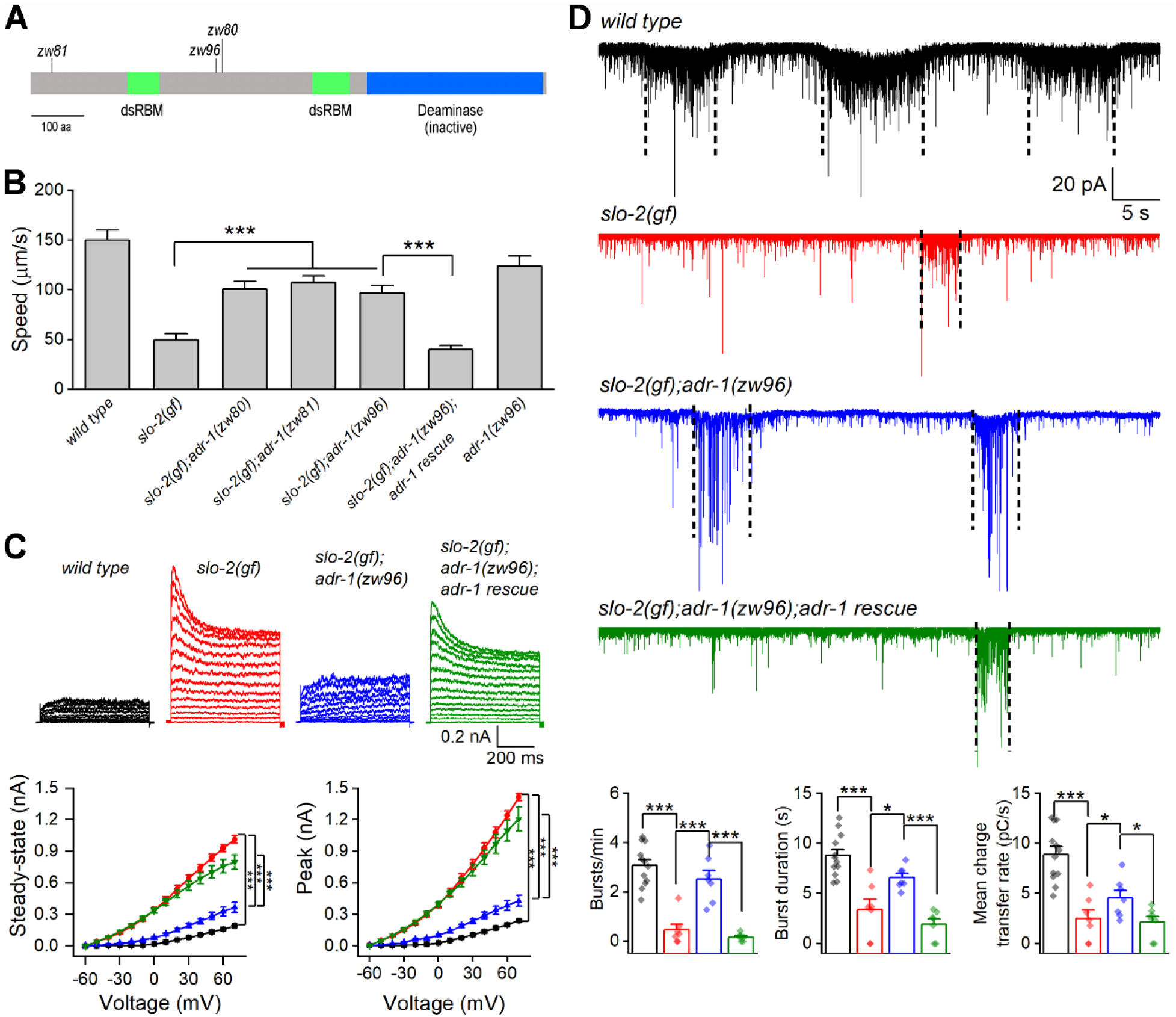
Loss-of-function mutations of *adr-1* suppress phenotypes caused by a hyperactive SLO-2. (**A**) Diagram of ADR-1 domain structures and locations of the non-sense mutations in the *adr-1* mutants. ADR-1 has two double-stranded RNA-binding motifs (dsRBM) and a pseudodeaminase domain. (**B**) Mutations of *adr-1* mitigated an inhibitory effect of hyperactive or gain-of-function (*gf*) SLO-2 on locomotion through acting in neurons. *adr-1* rescue was achieved by expressing GFP-tagged wild-type ADR-1 in neurons under the control of P*rab-3* (same in **C** and **D**). Sample sizes were 10–12 in each group. (**C**) *adr-1(zw96)* reduced an augmenting effect of *slo-2(gf)* on motor neuron whole-cell outward currents. Pipette solution I and bath solution I were used. Sample sizes were 14 in each group. (**D**) *adr-1(zw96)* mitigated an inhibitory effect of *slo-2(gf)* on postsynaptic current (PSC) bursts at the neuromuscular junction. The vertical dotted lines over the sample traces mark PSC bursts, which are defined as an apparent increase in PSC frequency accompanied by a sustained current (downward baseline shift) lasting > 3 sec. Pipette solution II and bath solution I were used. Sample sizes were 12 *wild type*, and 7 in each of the remaining groups. All values are shown as mean ± SE. The asterisks indicate statistically significant differences between indicated groups (\**p* < 0.05, \*\*\**p* < 0.001) based on either two-way (**C**) or one-way (D) ANOVA with Tukey’s post hoc tests. The following source data are available for Figure 1: **Source data 1.** Raw data and numerical values for data plotted in Figure 1.

In *C. elegans*, cholinergic motor neurons control body-wall muscle cells by producing bursts of postsynaptic currents (PSC bursts) (Liu et al., 2014). To determine how *adr-1* mutants might alleviate the *slo-2(gf)* locomotion defect, we recorded voltage-activated whole-cell currents from a representative cholinergic motor neuron (VA5) and postsynaptic currents from body-wall muscle cells in wild type, *slo-2(gf)*, *slo-2(gf);adr-1(zw96)*, and *slo-2(gf);adr-1(zw96)* with *adr-1* rescued in neurons. Compared with wild type, the *slo-2(gf)* strain displayed much larger outward currents, and greatly decreased PSC burst frequency, duration and charge transfer (**Fig. 1 C and D**). These phenotypes of *slo-2(gf)* were mostly suppressed in the *slo-2(gf);adr-1(zw96)* strain (**Fig. 1 C and D**), suggesting that *adr-1(lf)* alleviated the sluggish phenotype through inhibiting SLO-2(*gf*). In addition, expression of wild-type *adr-1* in neurons of *slo-2(gf);adr-1(zw96)* restored the effects of *slo-2(gf)* on whole-cell currents of VA5 and PSC bursts (**Fig. 1 C and D**). These observations suggest that inhibition of SLO-2 activity in motor neurons is likely a major contributor to the suppressing effect of *adr-1(lf)* on the *slo-2(gf)* sluggish phenotype.

We suspected that the suppression of SLO-2(*gf*) by *adr-1(lf)* resulted from deficient RNA-editing. If so, *adr-2(lf)* might similarly suppress the sluggish phenotype of *slo-2*(*gf*) as did *adr-1(lf)* because ADR-2 is required for RNA editing. Indeed, the sluggish phenotype of *slo-2(gf)* worms was substantially alleviated in *slo-2(gf);adr-2(lf)* double mutants (**Fig. 2A**). Also, the augmenting effect of *slo-2(gf)* on VA5 whole-cell outward currents was mostly abolished by *adr-2(lf)* (**Fig. 2B**). Furthermore, *adr-2(lf)* brought VA5 whole-cell currents below the wild-type level (**Fig. 2B**), which presumably resulted from reduced activities of wild-type SLO-2. These results suggest that RNA editing is important to SLO-2 function in neurons.

**Figure 2.**
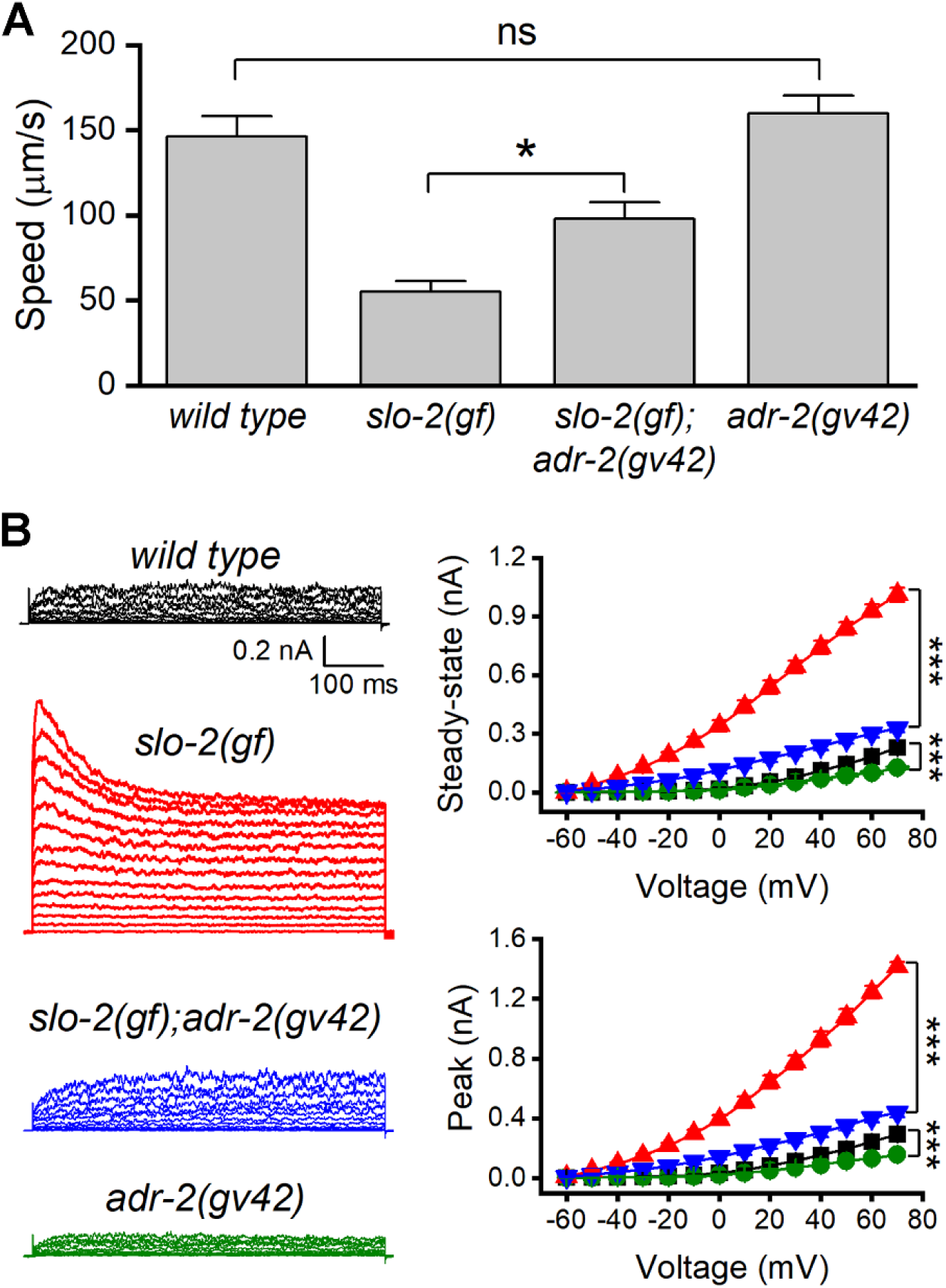
Loss-of-function mutation of *adr-2* suppressed the effects of gain-of-function (*gf*) *slo-*2 on locomotion and motor neuron whole-cell currents. (**A**) *adr-2(gv42)* alleviated an inhibitory effect of *slo-2(gf)* on locomotion speed. The sample size was 10–12 in each group. (**B**) *adr-2(gv42)* largely reversed an augmenting effect of *slo-2(gf)* on whole-cell currents in VA5 motor neuron. Sample sizes were 14 in each group. All data are shown as mean ± SE. Pipette solution I and bath solution I were used. The asterisks indicate statistically significant differences (* *p* < 0.05; *** *p* < 0.001) whereas “ns” stands for “no significant difference” between the indicated groups based on either one-way (**A**) or two-way (**B**) ANOVA with Tukey’s post hoc tests. The following source data are available for Figure 2: **Source data 1.** Raw data and numerical values for data plotted in Figure 2.

### ADR-1 is expressed in neurons and localized in the nucleus

The expression pattern of *adr-1* was examined by expressing GFP under the control of *adr-1* promoter (P*adr-1*). In transgenic worms, strong GFP expression was observed in the nervous system, including ventral cord motor neurons and many neurons in the head and tail, and weak GFP expression was observed in the intestine and body-wall muscles (**Fig. 3A**). We then examined the subcellular localization pattern of ADR-1 by expressing GFP-tagged full-length ADR-1 (ADR-1::GFP) under the control of P*rab-3*. We found that ADR-1::GFP is localized in the nucleus, as indicated by its colocalization with the mStrawberry-tagged nucleus marker HIS-58 (Liu et al., 2018) in ventral cord motor neurons (**Fig. 3B**).

**Figure 3.**
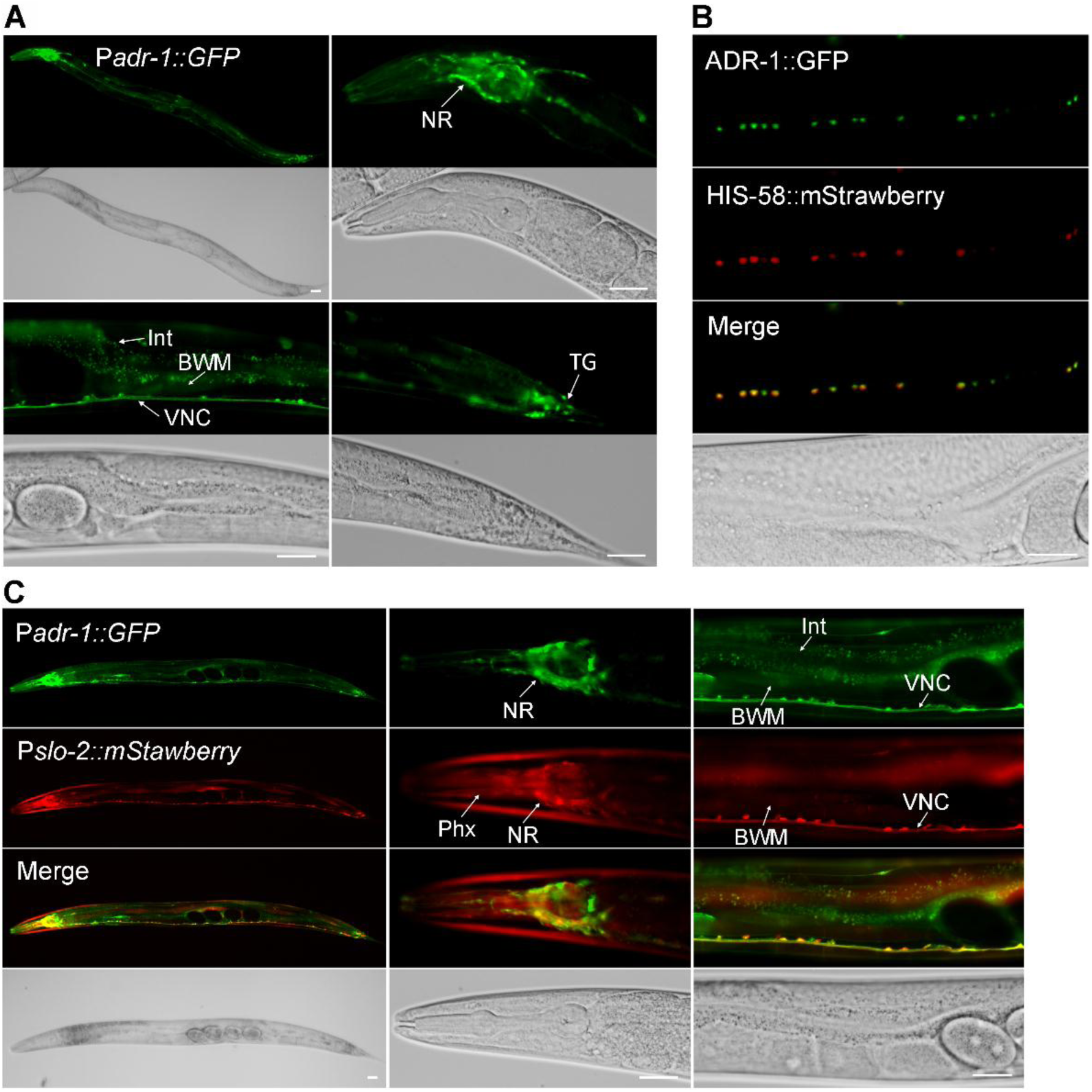
ADR-1 is coexpressed with SLO-2 in many neurons and localized in the nucleus. (**A**) Expression of an *adr-1* promoter (P*adr-1*)::GFP transcriptional fusion in worms resulted in strong GFP signal in many neurons (NR, nerve ring; VNC, ventral nerve cord; TG, tail ganglion) and weak GFP signal in body-wall muscles (BWM) and intestine (Int). (**B**) GFP-tagged ADR-1 (ADR-1::GFP) colocalized with a mStrawberry-tagged HIS-58 nucleus marker, as indicated by fluorescence images of VNC motor neurons. (**C**) *adr-1* and *slo-2* are co-expressed in many neurons but show differential expressions in the pharynx (Phx) and Int. Scale bar = 20 µm in in all panels.

To determine whether *adr-1* is co-expressed with *slo-2*, we crossed the P*adr-1::GFP* transgene into an existing strain expressing P*slo-2::*mStrawberry (Liu et al., 2018). We found that the expression patterns of *adr-1* and *slo-2* overlapped extensively in the nervous system (**Fig. 3C**). For example, the majority of ventral cord motor neurons and numerous head neurons were colabeled by GFP and mStrawberry (**Fig. 3C**). The occasional non-overlapping expressions of GFP and mStrawberry in ventral cord motor neurons probably resulted from mosaic expression of the transgenes.

### ADR-1 regulates neurotransmitter release through SLO-2

SLO-2 is the primary conductor of delayed outward currents in *C. elegans* cholinergic motor neurons (Liu et al., 2014). We wondered whether the function of native SLO-2 channels in motor neurons depends on ADR-1. Consistent with our previous report (Liu et al., 2014), VA5 delayed outward currents were dramatically smaller and VA5 resting membrane potential was much less hyperpolarized in *slo-2(lf)* than wild type. While *adr-1(lf)* also caused significantly decreased outward currents and less hyperpolarized resting membrane potential in VA5, it did not produce additive effects when combined with *slo-2(lf)* (**Fig. 4 A-C**). These results suggest that *adr-1(lf)* affects motor neuron outward current and resting membrane potential through SLO-2.

**Figure 4.**
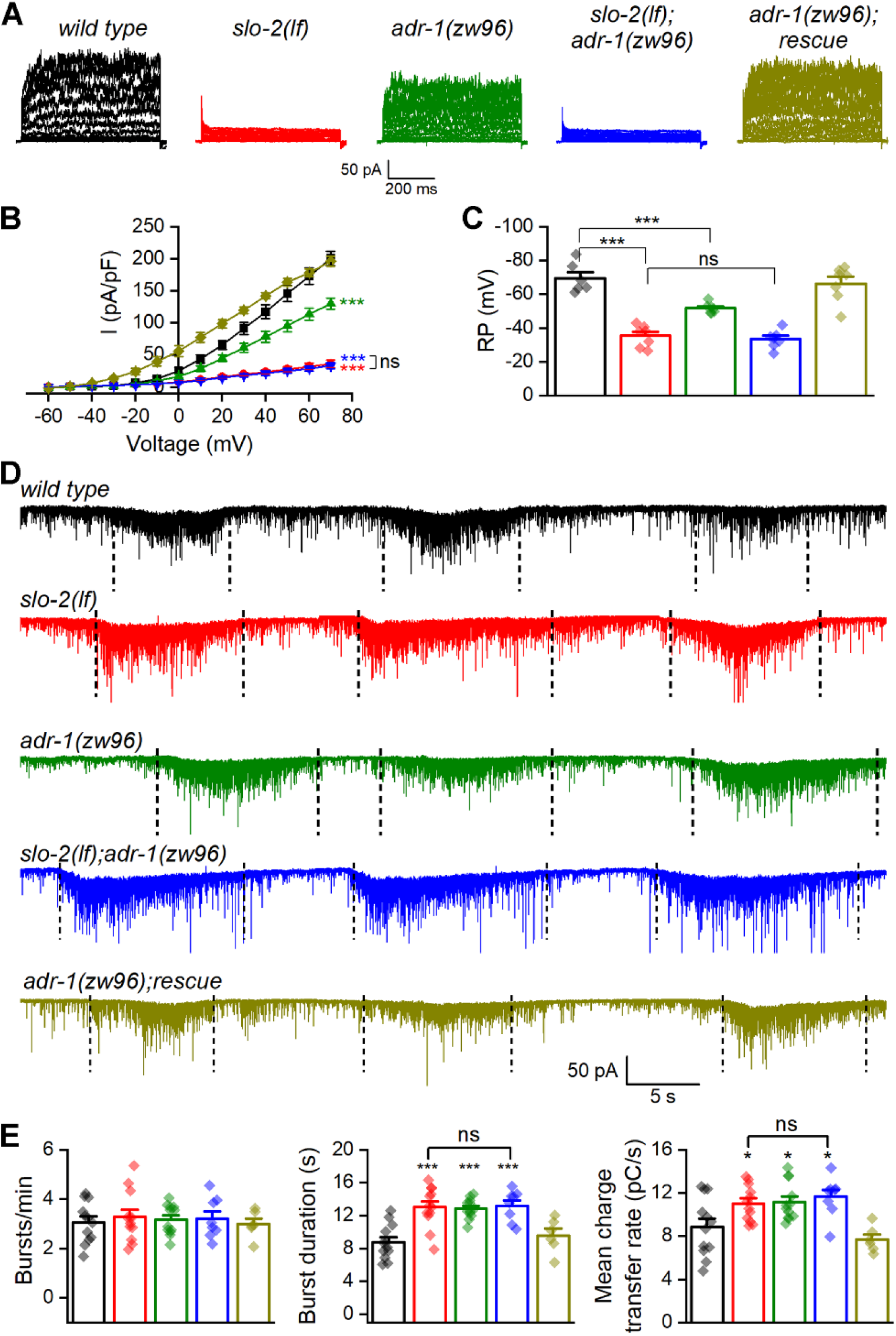
ADR-1 contributes to motor neuron whole-cell currents and regulates postsynaptic current (PSC) bursts through SLO-2. (**A**) Representative VA5 whole-cell current traces. (**B**) Current (*I*) - voltage relationships of the whole-cell currents. Sample sizes were 14 in each group. (**C**) Resting membrane potentials of VA5. Sample sizes were 6 *wild type*, and 7 in each of the remaining groups. (**D**) Representative traces of spontaneous PSCs with PSC bursts marked by vertical dotted lines. (**E**) Comparisons of PSC burst properties. Sample sizes were 8 *slo-2(nf101);adr-1(zw96),* 6 *adr-1(zw96)* rescue, and 12 in each of the remaining groups. All values are shown as mean ± SE. The asterisks indicate statistically significant differences (\**p* < 0.05, \*\*\**p* < 0.001) compared with *wild type* whereas “ns” stands for no significant difference between the indicated groups based on either two-way (**B**) or one-way (**C** and **E**) ANOVA with Tukey’s post hoc tests. Pipette solution I and bath solution I were used in (**A**) and (**C**). Pipette solution II and bath solution I were used in (**D**). The following figure supplement and source data are available for Figure 4:

We next determined whether *adr-1(lf)* also alters PSC bursts. We found that *adr-1(lf)* caused an increase in the duration and mean charge transfer rate of PSC bursts without altering the burst frequency compared with wild type (**Fig. 4 D and E**). These phenotypes of *adr-1(lf)* were similar to those of *slo-2(lf)* and did not become more severe in the double mutants (**Fig. 4 D and E**), suggesting that ADR-1 modulates neurotransmitter release through SLO-2. The similar effects of *adr-1(lf)* and *slo-2(lf)* on PSC bursts are in contrast to their differential effects on VA5 outward currents and resting membrane potential. This discrepancy suggests that there might be a threshold level of SLO-2 deficiency to cause a similar change in PSC bursts.

### ADR-1 regulates SLO-2 function through SCYL-1

Given that our results suggest that RNA editing is important to SLO-2 function, we determined whether *adr-1(lf)* causes deficient editing or decreased expression of *slo-2* mRNA by comparing RNA-seq data between *adr-1(lf)* and wild type. The *adr-1(zw96)* allele was chosen for these analyses to minimize potential complications by mutations of other genes introduced in *adr-1* mutants isolated from the genetic screen. Unexpectedly, no RNA editing event was detected in *slo-2* transcripts, and *slo-2* mRNA level was similar between wild type and the *adr-1* mutant (**Fig. 4-figure supplement 1**). These results suggest that ADR-1 might regulate SLO-2 function through RNA editing of another gene.

A previous study identified 270 high-confidence editing sites in transcripts of 51 genes expressed in *C. elegans* neurons (Washburn et al., 2014). We suspected that the putative molecule mediating the effect of ADR-1 on SLO-2 might be encoded by one of these genes, and the mRNA level of this gene may have reduced expression in *adr-1(lf)*. Therefore, we compared transcript expression levels of these genes (excluding those encoding transposons) quantified from our RNA-Seq data between wild type and *adr-1(zw96)*. The transcripts of most genes showed either no decrease or only a small decrease, but two of these genes, *rncs-1* and *scyl-1*, were reduced greatly in *adr-1(lf)* compared with wild type (**Fig. 5**). *rncs-1* is not a conceivable candidate for the putative SLO-2 regulator because it is a non-coding gene expressed in the hypodermis and vulva (Hellwig and Bass, 2008). On the other hand, *scyl-1* is a promising candidate because it encodes an orthologue of mammalian SCYL1 important to neuronal function and survival (Pelletier, 2016). We therefore focused our analyses on *scyl-1*. Like its mammalian homologs, SCYL-1 has an amino-terminal kinase domain that lacks residues critical to kinase activity, and a central domain containing five HEAT repeats (HEAT for *H*untingtin, *e*longation factor 3, protein phosphatase 2*A*, yeast kinase *T*OR1) (Pelletier, 2016). SCYL-1 shares 38% identity and 60% similarity with human SCYL1. Notably, amino acid sequence in the HEAT domain, which is often highly degenerative (Pelletier, 2016), shows a very high level of sequence homology (53% identity and 76% similarity) between these two proteins (**Fig. 5-figure supplement 1**).

**Figure 5.**
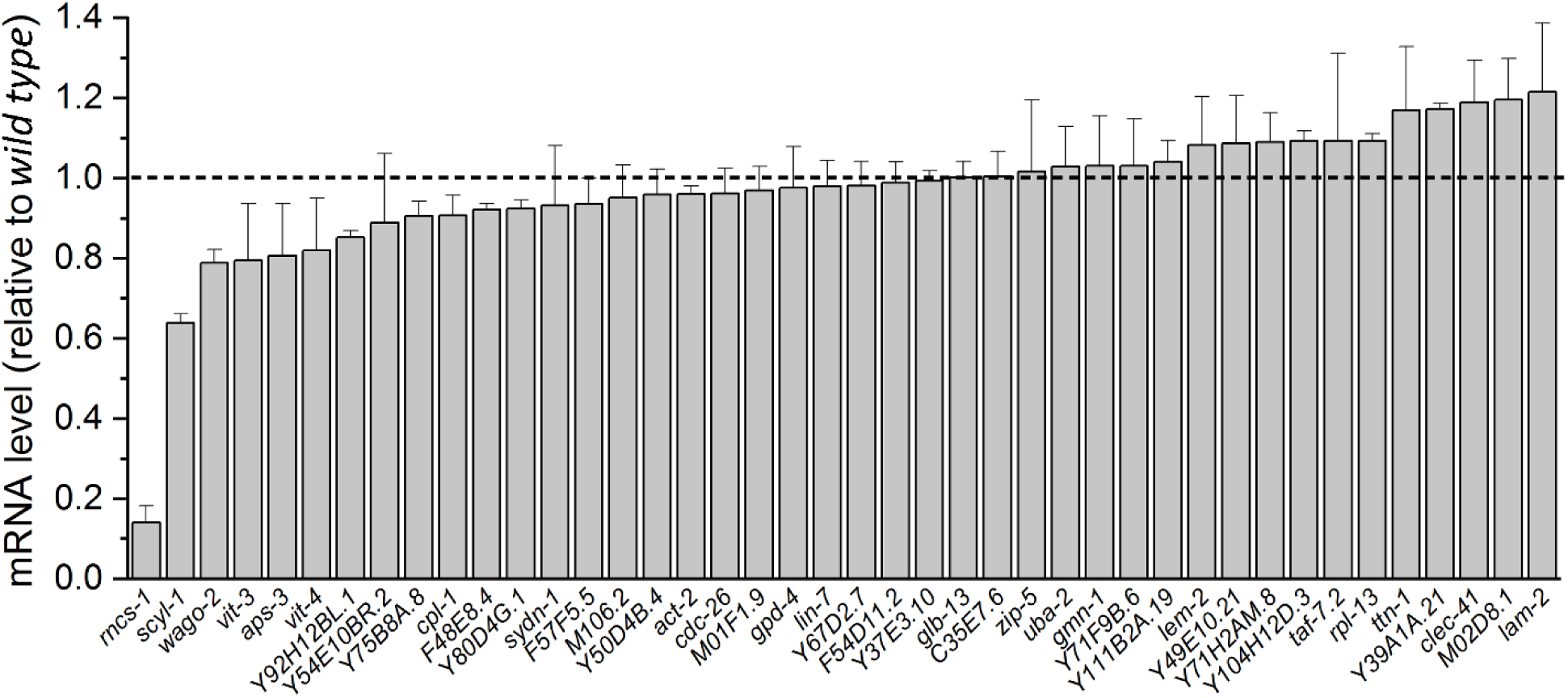
Normalized transcript expression levels of selected genes in *adr-1(zw96)* mutant. The genes were selected based on the detection of ADR-1-dependent RNA editing events in their transcripts reported in an earlier study (Washburn et al., 2014). Transcript expression level of each gene in the mutant is normalized by that in the wild type. Shown are mean ± SE from three biological replicates of RNA-seq experiments. The following figure supplement and source data are available for Figure 5:

We first examined the expression pattern of *scyl-1* by expressing GFP reporter under the control of *scyl-1* promoter (P*scyl-1*). An *in vivo* homologous recombination approach was used in this experiment to include a large fragment of genomic DNA sequence upstream of the *scyl-1* initiation site. Specifically, a 0.5-kb genomic fragment upstream of the *scyl-1* initiation site was cloned by PCR and fused to GFP. The resultant plasmid was co-injected with a fosmid covering part of the *scyl-1* coding region and 32 kb sequence upstream of the initiation site into wild type worms. *In vivo* homologous recombination between the plasmid and the fosmid is expected to result in a P*scyl-1*::GFP transcriptional fusion that includes all the upstream sequence in the fosmid. After successful creation of a transgenic strain expressing the P*scyl-1*::GFP transcriptional fusion, we crossed the transgene into the P*slo-2*::mStrawberry strain, and examined the expression patterns of GFP and mStrawberry. We observed co-expression of *scyl-1* and *slo-2* in many ventral cord motor neurons (**Fig. 6**). However, most other neurons expressing *slo-2* (*e. g.* head and tail neurons) did not appear to express *scyl-1*. In addition, *scyl-1* expression was detected in some cells that did not express *slo-2*, including the excretory cell, spermatheca, vulval muscle cells, and intestinal cells (**Fig. 6**).

**Figure 6.**
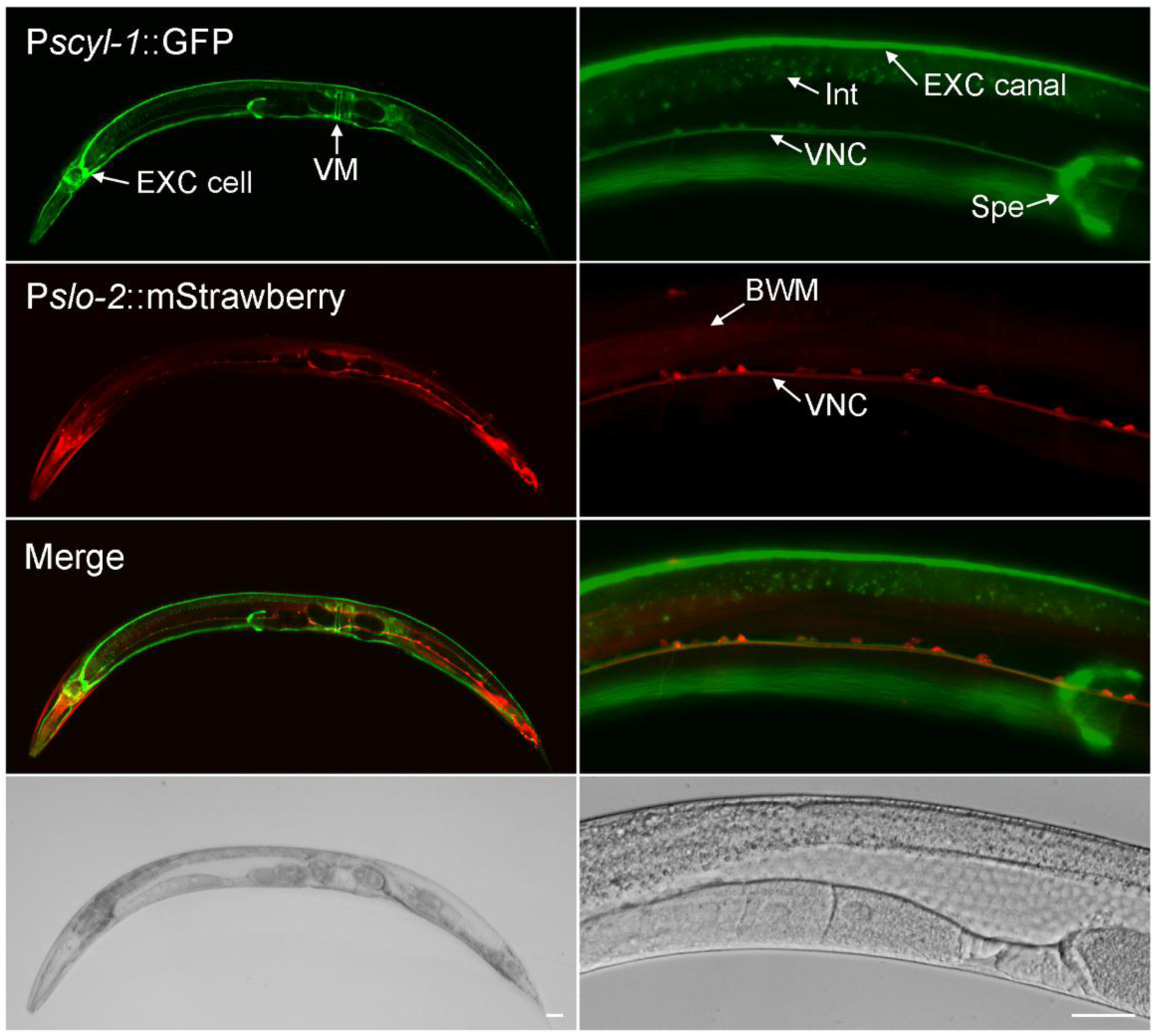
*scyl-1* and *slo-2* are coexpressed in ventral cord motor neurons but differentially expressed in other cells. In transgenic worms coexpressing P*scyl-1::GFP* and P*slo-2::mStrawberry* transcriptional fusions, GFP signal was observed in ventral nerve cord (VNC) motor neurons, the large H-shaped excretory (EXC) cell, vulval muscles (VM), and spermatheca (Spe) while mStrawberry signal was detected in VNC motor neurons, body-wall muscles (BMW), and many other neurons. Scale bar = 20 µm.

We next determined whether SCYL-1 is related to SLO-2 function. To this end, we created a mutant, *scyl-1(zw99)*, by introducing a stop codon after isoleucine 152 using the CRISPR/Cas9 approach, and examined the effect of this mutation on VA5 delayed outward currents. *scyl-1(zw99)* showed a substantial decrease in VA5 outward currents compared with wild type; and this phenotype was non-additive with that of *slo-2(lf)* and could be rescued by expressing wild type SCYL-1 in neurons (**Fig. 7A**). These results suggest that SCYL-1 contributes to SLO-2-dependent outward currents.

**Figure 7.**
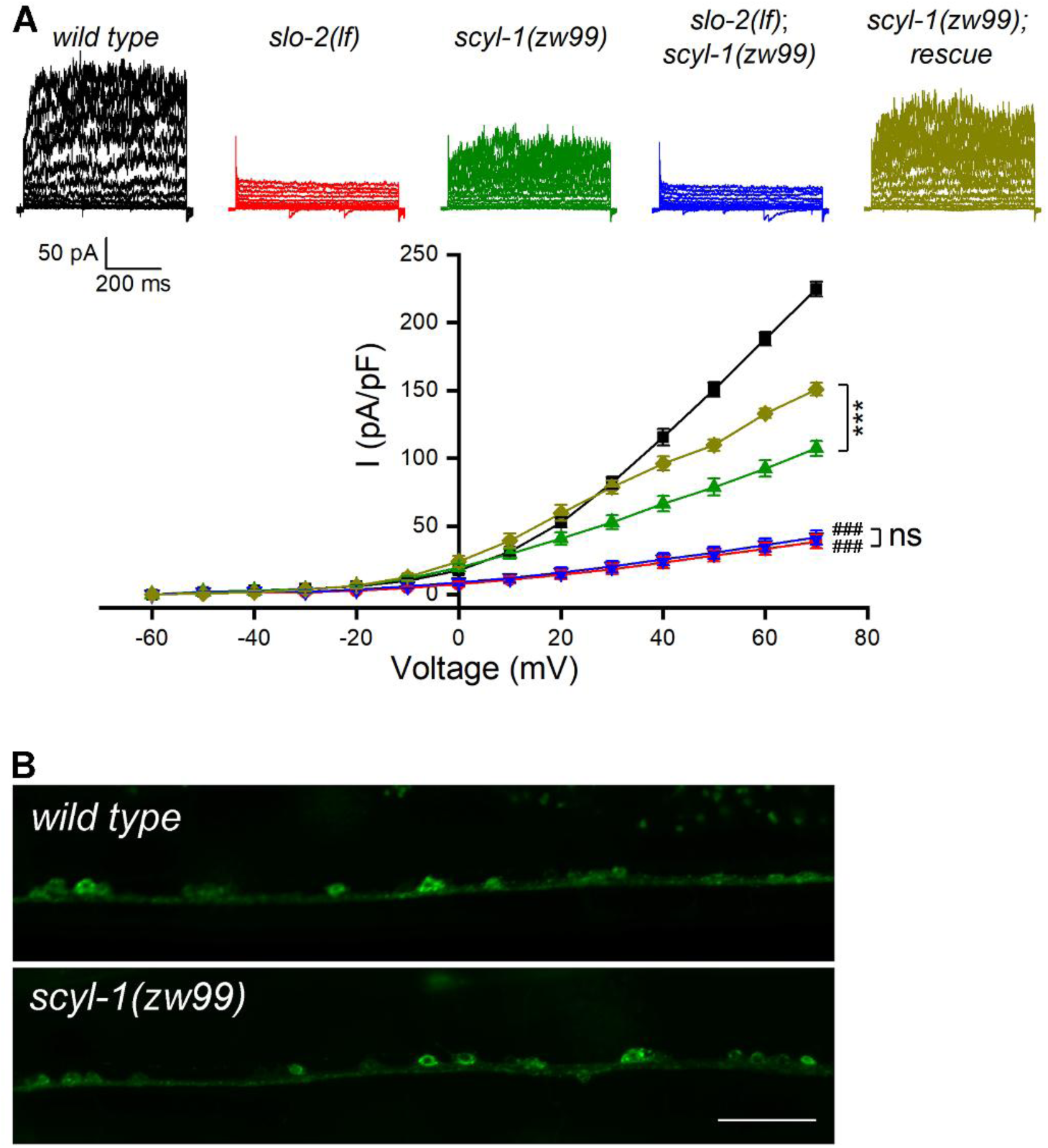
SCYL-1 contributes to motor neuron outward currents through SLO-2. (**A**) Sample whole-cell current traces of VA5 motor neurons and the current-voltage relationships. Sample sizes were 14 in each group. The rescue strain was created by expressing wild-type *scyl-1* under the control of P*rab-3*. All values are shown as mean ± SE. The asterisks (***) and pound signs (^###^) indicate statistically significant differences (*p* < 0.001) between the indicated groups and from wild type, respectively, whereas “ns” stands for no significant difference between the indicated groups (two-way ANOVA with Tukey’s post hoc tests). (**B**) GFP signal in ventral cord motor neurons was indistinguishable between *wild type* and *scyl-1(zw99)* worms expressing GFP-tagged full-length SLO-2 under the control of P*rab-3*. Scale bar = 20 µm. The following source data are available for Figure 7: **Source data 1.** Raw data and numerical values for data plotted in Figure 7.

The decrease of delayed outward currents in *scyl-1(lf)* could have resulted from decreased expression or function of SLO-2. We first determined whether *scyl-1(lf)* alters SLO-2 expression by crossing a stable (near 100% penetrance) P*rab-3*::SLO-2::GFP transgene from an existing transgenic strain of wild-type genetic background (Liu et al., 2018) into *scyl-1(zw99)*, and comparing GFP signal between the two strains. We found that GFP signal in the ventral nerve cord was similar between wild type and the *scyl-1* mutant (**Fig. 7B**), suggesting that SCYL-1 does not regulate SLO-2 expression. We then determined whether SCYL-1 regulates SLO-2 function by obtaining inside-out patches from VA5 and analyzing SLO-2 single-channel properties. SLO-2 showed >50% decrease in open probability (*P_o_*) without a change of single-channel conductance in *scyl-1(zw99)* compared with wild type, and this mutant phenotype was completely rescued by neuronal expression of wild-type SCYL-1 (**Fig. 8A**). Analyses of single-channel open and closed events revealed that SLO-2 has two open states and three closed states, and that the decreased *P_o_* of SLO-2 in *scyl-1(lf)* mainly resulted from decreased events of long openings (**Fig. 8B**) and increased events of long closures (**Fig. 8C**).

**Figure 8.**
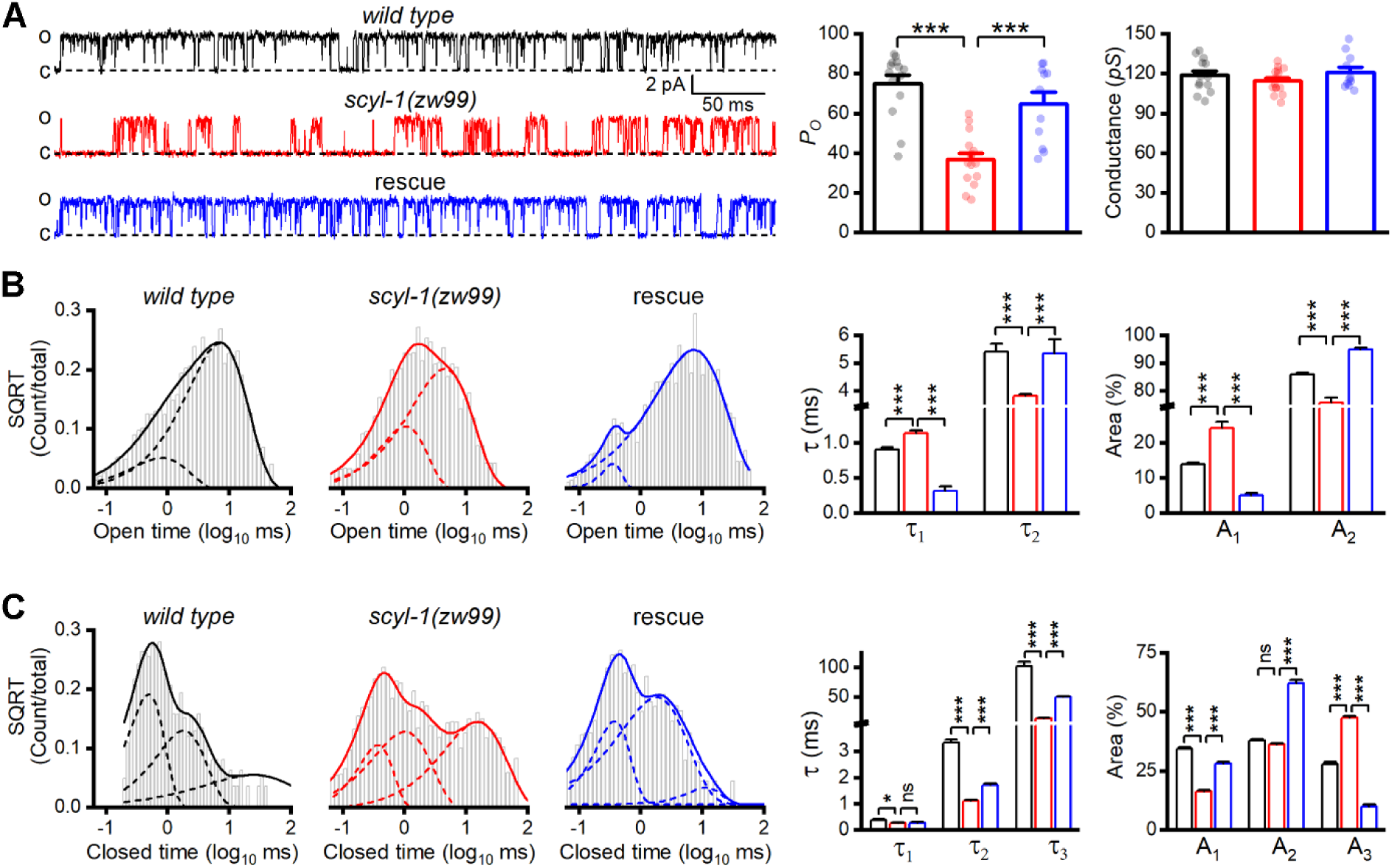
Single-channel open probability (P_o_) of SLO-2 is decreased in *scyl-1* mutant. (**A**) Representative SLO-2 single-channel currents from inside-out patches of the VA5 motor neuron, and comparisons of *P_o_* and single-channel amplitude between *wild type* (*n* = 14), *scyl-1(zw99)* (*n* = 15), and *scyl-1(zw99)* rescued by expressing wild-type *scyl-1* in neurons under the control of P*rab-3* (*n* = 11). (**B** and **C**) Fitting of open (**B**) and closed (**C**) durations to exponentials, and comparisons of τ values and relative areas of the fitted components (indicated by dotted lines). Pipette solution III and bath solution II were used. All values are shown as mean ± SE. The asterisks indicate a significant difference between the indicated groups (* *p* < 0.05, *** *p* < 0.001, one-way ANOVA with Tukey’s post hoc tests). The following source data are available for Figure 8: **Source data 1.** Raw data and numerical values for data plotted in Figure 8.

The observed effects of *scyl-1(lf)* on SLO-2 single-channel properties suggest that SCYL-1 may physically interacts with SLO-2. We performed bimolecular fluorescence complementation (BiFC) assays (Hu et al., 2002) to test this possibility. In these assays, full-length SCYL-1 tagged with the carboxyl terminal portion of YFP (YFPc) was coexpressed in neurons with either full-length, amino terminal portion, or carboxyl terminal portion of SLO-2 tagged with the amino terminal portion of YFP (YFPa) (**Fig. 9A**). YFP fluorescence was observed in ventral cord motor neurons when either the full-length or the C-terminal portion of SLO-2 was used but not when the N-terminal protein was used in the assays (**Fig. 9B**). These results suggest that SCYL-1 physically interacts with SLO-2, and this interaction depends on SLO-2 carboxyl terminal portion.

**Figure 9.**
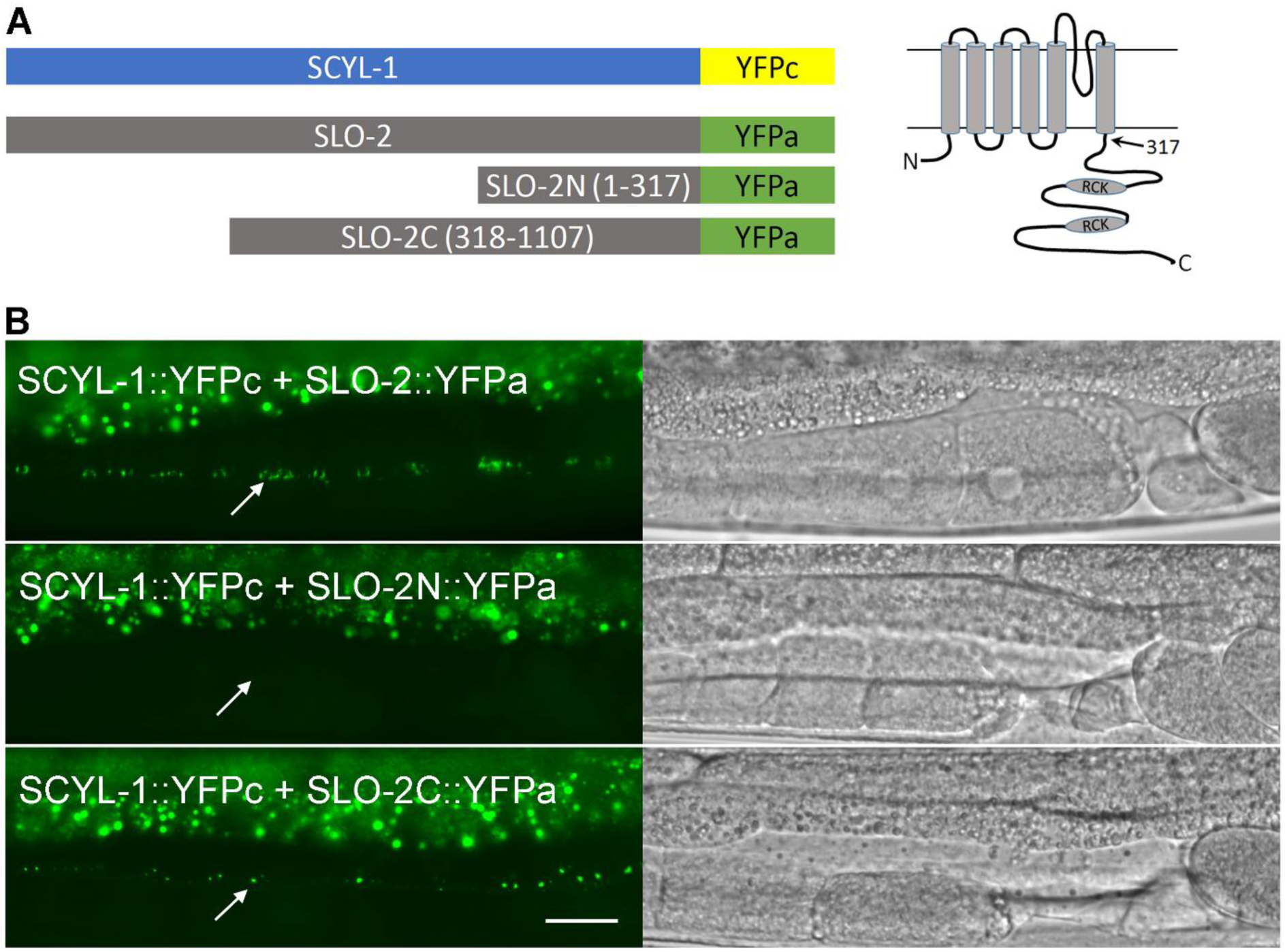
SCYL-1 physically interacts with SLO-2 in neurons. (**A**) Diagrams of the various fusion proteins used in the BiFC assays (*left*) and of SLO-2 membrane topology (*right*). The arrow indicates the split site for SLO-2N and SLO-2C fusions. RCK, regulator of conductance for K^+^. (**B**) YFP signal was detected when SCYL-1 was coexpressed with either full-length or the carboxyl terminal portion of SLO-2 but not with the amino terminal portion of SLO-2. Shown are representative fluorescent images of the ventral nerve cord (indicated by arrows) with corresponding DIC images. The bright signals at the top of each fluorescence image was from auto-fluorescence of the intestine. Scale bar = 20 μm.

### scyl-1 expression depends on RNA editing at a specific 3’-UTR site

Our RNA-Seq data revealed eight high-frequency (>15%) adenosine-to-guanosine editing sites in *scyl-1* transcripts of wild type (**Fig. 10A**). All these editing sites are located within a predicted 746 bp hair-pin structure in the 3’ end of *scyl-1* pre-mRNA, which contains an inverted repeat with >98% complementary base pairing (**Fig. 10B**). Interestingly, RNA editing at only one of the eight sites was significantly undermined (by 74%) in *adr-1(zw96)* compared with wild type (**Fig. 10A**). Sanger sequencing of *scyl-1* mRNA and the corresponding genomic DNA from wild type, *adr-1(zw96)*, and *adr-2(gv42)* confirmed that RNA editing at this specific site was deficient in both the *adr-1* and *adr-2* mutants whereas editing at an adjacent site was deficient only in the *adr-2* mutant (**Fig. 10C**), suggesting that RNA editing at the site impaired by *adr-1(lf)* might be important to *scyl-1* expression. To test this possibility, we fused GFP coding sequence in-frame to a genomic DNA fragment covering part of the last exon of *scyl-1* and 5 kb downstream sequence, and expressed it in neurons under the control of P*rab-3* (**Fig. 10D**). We also made a modified plasmid construct in which adenosine (A) was changed to guanosine (G) at the specific ADR-1-dependent editing site to mimic the editing (**Fig. 10D**). In transgenic worms harboring the original genomic sequence, no GFP signal was detected in neurons (**Fig. 10E**). In contrast, strong GFP signal was observed in neurons of transgenic worms expressing the A-to-G mutated genomic sequence (**Fig. 10E**). Taken together, the results suggest that ADR-1 plays a key role in *scyl-1* expression by promoting RNA editing at a specific site in its 3’-UTR.

**Figure 10.**
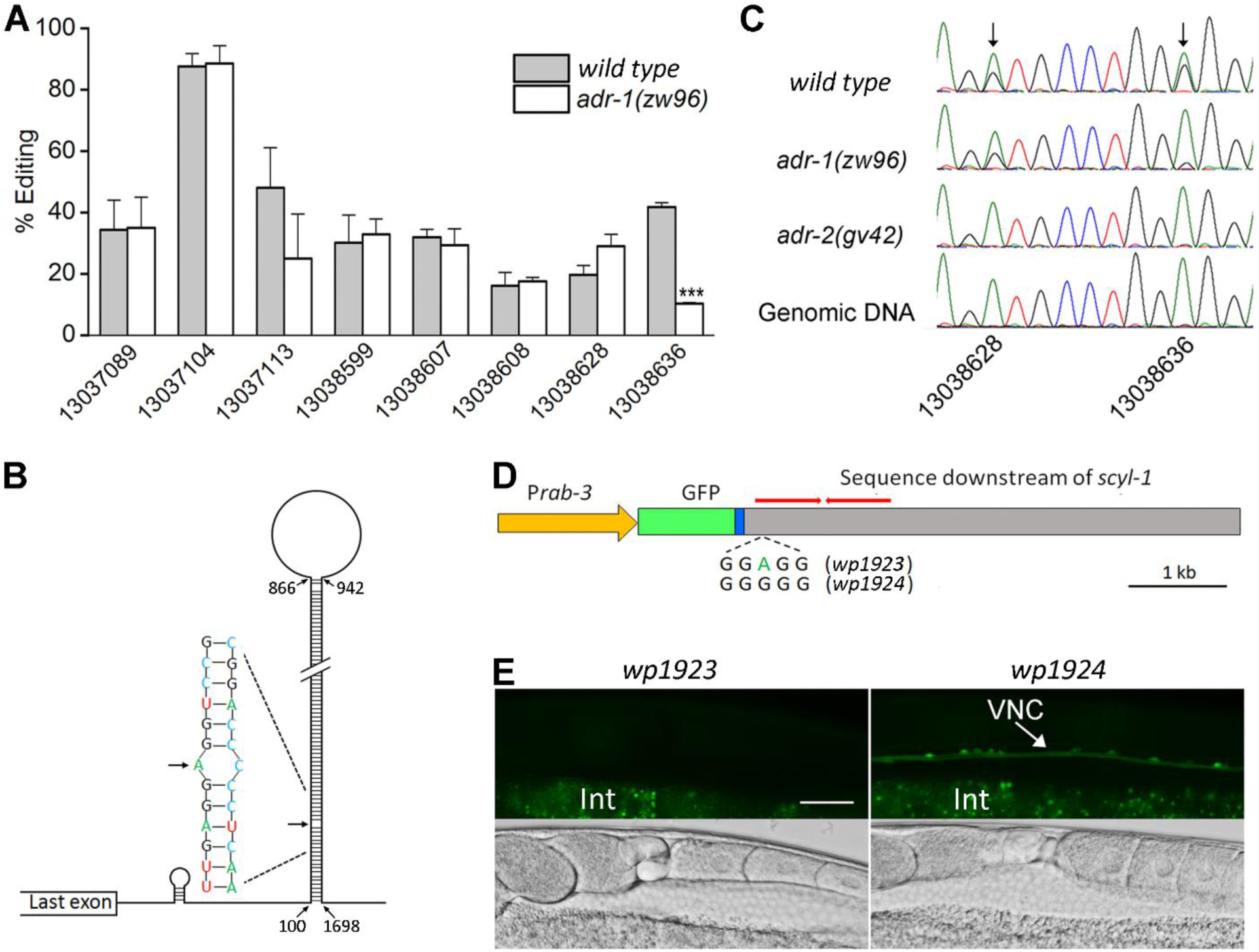
ADR-1 regulates *scyl-1* expression through RNA editing at a specific nucleotide in the 3’-UTR. (**A**) RNA editing at one out of eight highly (>15%) edited sites is severely deficient in *adr-1(zw96)* compared *wild type*. The percentage of editing was calculated by diving the number of reads containing A-I conversion by the total number of reads at each site. The *x*-axis indicates the positions of the edited adenosines in chromosome *V* (NC_003283). Shown are results (mean ± SE) of three RNA-seq experiments. The asterisks (***) indicate a statistically significant difference (*p* < 0.001, unpaired *t*-test). (**B**) Diagram showing a predicted hair-pin structure in the 3’end of *scyl-1* pre-mRNA with 746 complementary base pairs. Nucleotide are numbered from the first nucleotide of the 3’-UTR. (**C**) Chromatograms of *scyl-1* mRNA 3’-UTRs of *wild type*, *adr-1(zw96)*, and *adr-2(gv42)*, and of the corresponding *wild type* genomic DNA. Two editing sites in *wild type* mRNA (indicated by arrows) display a mixture of green (adenosine) and black (guanosine) peaks. While both editing events are non-existent in *adr-2(gv42)*, only one of them is inhibited by *adr-1(zw96)*. (**D**) Diagram of two GFP reporter constructs (*wp1923* and *wp1924*) used to confirm the role of the ADR-1-dependent editing site in gene expression. GFP was placed under the control of P*rab-3* and fused to the last exon (blue) of *scyl-1* followed by 5 kb downstream genomic sequence. The red bars indicate the inverted repeat sequences that form the double-stranded RNA in the hair-pin structure (**B**). *wp1923* contains the intact genomic sequence of *scyl-1* 3’-UTR, whereas *wp1924* differs from it in an A-to-G conversion mimicking the ADR-1-dependent editing. (**E**) Effects of the A-to-G conversion on GFP reporter expression. Shown are fluorescent and corresponding DIC images of transgenic worms harboring either *wp1923* or *wp1924*. GFP expression in the ventral nerve cord (VNC) was observed only in worms harboring *wp*1924. The diffused signal at the bottom of each fluorescent image was from auto-fluorescence of the intestine (Int). Scale bar = 20 µm. The following source data are available for Figure 10: **Source data 1.** Raw data and numerical values for data plotted in Figure 10.

### Human Slo2.2/Slack is regulated by SCYL1

The HEAT domain of SCYL proteins play important roles in protein-protein interactions but generally varies considerably in amino acid sequence for interactions with different partners (Yoshimura and Hirano, 2016). The high level of sequence homology of the HEAT domain between mammalian SCYL1 and worm SCYL-1 (**Fig. 5-figure supplement 1**) promoted us to test whether mammalian Slo2.2/Slack is also regulated by SCYL1. We expressed human Slack either alone or together with mouse SCYL1 in *Xenopus* oocytes, and analyzed Slack single-channel properties. Coexpression of SCYL1 caused ∼130% increase in Slack *P_o_* (**Fig. 11A**). The channel has at least two open states and two closed states. SCYL1 increased the duration and proportion of the long open state; and decreased the proportion but increased the duration of the long closed state (**Fig. 11 B and C**). These effects of SCYL1 on Slack are similar to those of SCYL-1 on SLO-2 single-channel properties (**Fig. 8**), suggesting that regulation of Slo2 channel function is likely a conserved physiological function of SCYL-1/SCYL1 proteins.

**Figure 11.**
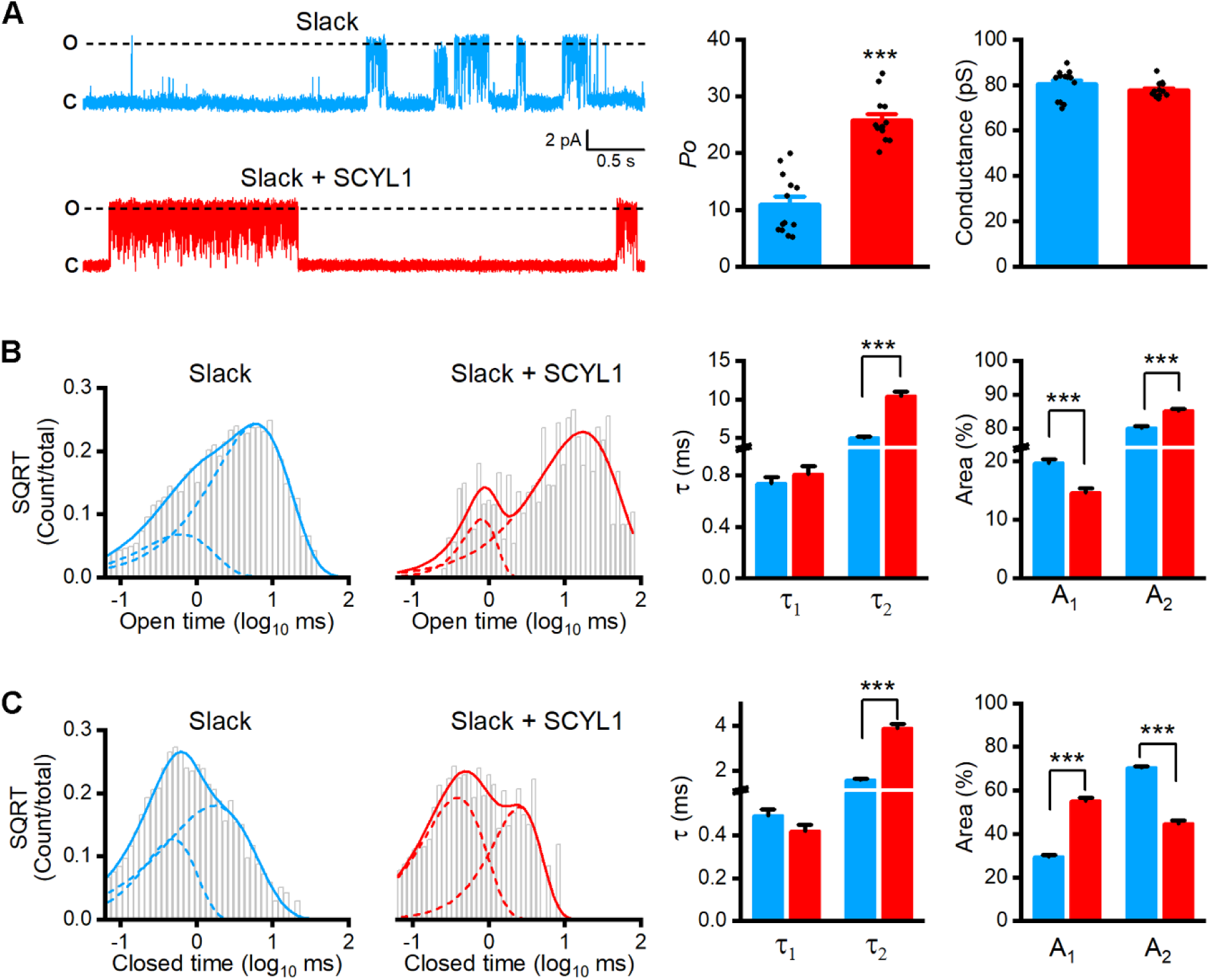
Single-channel open probability (*P_o_*) of human Slo2.2/Slack is augmented by SCYL1 in *Xenopus* oocyte expression system. (**A**) Representative traces of single-channel currents from inside-out patches and comparisons of *P_o_* and single-channel amplitude between patches with and without mouse SCYL1. (**B** and **C**) Fitting of open (**B**) and closed (**C**) durations to exponentials, and comparisons of τ values and relative areas of the fitted components (indicated by dotted lines) between the two groups. Sample sizes were 13 in both groups. All values are shown as mean ± SE. The asterisks (***) indicate a significant difference compared between the indicated groups (*p* < 0.001, unpaired *t*-test). The following source data are available for Figure 11: **Source data 1.** Raw data and numerical values for data plotted in Figure 11.

## Discussion

This study shows that both ADR-1 and SCYL-1 are critical to SLO-2 physiological function in neurons. While ADR-1 enhances SLO-2 function indirectly through regulating the expression level of SCYL-1, the latter do so directly. These conclusions are supported by multiple lines of evidence, including the isolation of *adr-1(lf)* mutants as suppressors of SLO-2(*gf*), the inhibition of SLO-2 activities by either *adr-1(lf)* or *scyl-1(lf)*, the reduction of *scyl-1* transcript expression in *adr-1(lf)* and correlation between *scyl-1* RNA editing and gene expression, the SLO-2 carboxyl terminal-dependent reconstitution of YFP fluorophore in BiFC assays with SCYL-1, and the inhibitory effects of *scyl-1(lf)* on SLO-2 single-channel activities. Importantly, we found that the human Slack is also regulated by SCYL1.

The biological significance of RNA editing at non-coding regions is only beginning to be appreciated. A recent study with *C. elegans* identified many neural-specific A-to-I editing sites in the 3’-UTR of *clec-41*, and found that *adr-2(lf)* causes both an elimination of these editing events and a chemotaxis defect (Deffit et al., 2017). Although it is unclear how *clec-41* expression is controlled by these editing events, and a direct link between the chemotaxis defect of *adr-2(lf)* mutant and the decreased *clec-41* expression remains to be established, these results suggest that RNA editing at non-coding regions might have important biological functions. In the present study, we demonstrate that A-to-I RNA editing at the 3’UTR of *scyl-1* controls its expression, and that SCYL-1 contributes to neuronal whole-cell currents through a direct effect on the SLO-2 channel. The results of these two studies have provided a glimpse of the biological roles of 3’-UTR RNA editing in gene expression and neuronal function.

Our results demonstrate that RNA editing at a single site in the 3’-UTR could have a profound effect in promoting gene expression. The A-to-I conversion at the specific editing site of *scyl-1* increases base pairing in the putative double-stranded structure of the 3’-UTR (**Fig. 10B**). Increased base paring in a double-stranded RNA generally facilitates RNA degradation. It is therefore intriguing how such an increased paring in the 3’-UTR may cause increased gene expression. One possibility is that editing at this site helps recruit a specific RNA-binding protein to the 3’-UTR to prevent *scyl-1* mRNA from degradation. Although the exact mechanism remains to be determined, it is a remarkable first example that a specific RNA editing site at the 3’UTR plays a crucial role in gene expression.

SCYL1 proteins are evolutionarily conserved proteins that share an N-terminal pseudokinase domain (Manning et al., 2002; Pelletier, 2016). Results of previous studies with cultured cells suggest that SCYL1 may regulate intracellular trafficking processes between the Golgi apparatus and the ER (Burman et al., 2008; Burman et al., 2010), and facilitate nuclear tRNA export by acting at the nuclear pore complex (Chafe and Mangroo, 2010). Mutations of SCYL1 in humans are associated with a variety of disorders, including neurodegeneration, intellectual disabilities, and liver failure (Lenz et al., 2018; Li et al., 2019; Schmidt et al., 2015; Shohet et al., 2019; Spagnoli et al., 2018, 2019). Mice with SCYL1 deficiency show an early onset and progressive neurodegenerative disorder (Pelletier et al., 2012). However, it is unclear whether the documented mutant phenotypes of SCYL1 are related to its known roles in intracellular trafficking and nuclear tRNA export (Pelletier, 2016). The results of this study indicate a new role of SCYL1/SCYL-1 proteins: regulating Slo2 channels. What might be the molecular mechanism through which SCYL-1 enhances SLO-2 activity? Since SCYL-1 physically associates with SLO-2, and enhances SLO-2 single-channel *P_o_* through altering the open and closed states, it likely regulates channel function either directly or through a closely associated protein. The fact that human Slack *P_o_* is augmented by mouse SCYL1 through effects on the open and closed states lends further support to the notion that SCYL1/SCYL-1 may regulate Slo2 channel activities through close interactions. While the exact mechanism remains to be determined, the observed large effects of SCYL-1/SCYL1 proteins on channel activities suggest that they are likely a major player in SLO-2/Slo2 physiological function.

The expression patterns of *scyl-1* and *slo-2* largely do not overlap. Although they are coexpressed in ventral cord motor neurons, where SCYL-1 is required for SLO-2 physiological function, most other neurons expressing *slo-2* do not express *scyl-1*, suggesting that the regulatory effect of SCYL-1 on SLO-2 is cell- and tissue-specific. Furthermore, *scyl-1* expression was observed in a variety of cells that do not express *slo-2*, suggesting that SCYL-1 physiological functions are not limited to regulating SLO-2. In mouse, SCYL1 and Slack are both expressed in the hippocampus and cerebellum, but their reported expression patterns do not completely overlap (Joiner et al., 1998; Schmidt et al., 2007). Conceivably, Slack channels might also exhibit cell- and tissue-specific dependence on SCYL1, and SCYL1 proteins likely perform other functions besides regulating Slack channels. The pleiotropic phenotypes associated with mutations of SCYL1 (Lenz et al., 2018; Li et al., 2019; Pelletier et al., 2012; Schmidt et al., 2015; Shohet et al., 2019; Spagnoli et al., 2018, 2019) support the notion that SCYL1 proteins are also important to other physiological functions.

In summary, this study demonstrates that ADAR-mediated RNA editing controls the expression of SCYL-1, which interacts with SLO-2 to allow SLO-2 perform its physiological functions. Moreover, this study shows that this regulatory mechanism is conserved with mammalian SCYL1 and Slo2. Our findings reveal a new molecular mechanism of Slo2 channel regulation, and provide the bases for investigating how Slo2 physiological functions are regulated by SCYL1, and whether the neurodegeneration and intellectual disability phenotypes of SCYL1 mutations are related to Slo2 channel dysfunction.

## Materials and Methods

### C. elegans culture and strains

*C. elegans* hermaphrodites were grown on nematode growth medium (NGM) plates spotted with a layer of OP50 *Escherichia coli* at 22°C inside an environmental chamber.The following strains were used in this study (plasmids used in making the transgenic strains are indicated by numbers with a “*wp*” prefix): wild type (Bristol N2). LY101: *slo-2(nf101)*. ZW860: *zwIs139*[*Pslo-1::slo-2(gf)(wp1311), Pmyo-2::YFP(wp214)*]. ZW876: *zwIs139*[*Pslo-1::slo-2(gf)(wp1311), Pmyo-2::YFP(wp214)*]*; adr-1(zw80)*. ZW877: *zwIs139*[*Pslo-1::slo-2(gf)(wp1311), Pmyo-2::YFP(wp214)*]*; adr-1(zw81)*. ZW983: *zwIs139*[*Pslo-1::slo-2(gf)(wp1311), Pmyo-2::YFP(wp214)*]*; adr-2(gv42)*. ZW1002: *adr-2(gv42)*. ZW1049: *zwEx221*[*Prab-3::slo-2::GFP*]. ZW1388: *zwEx260*[*Prab-3::His-58::mStrawberry(p1749), Prab-3::adr-1::GFP(p1374)*]. ZW1394: *adr-1(zw96)*. ZW1401: *zwEx261*[*Padr-1::GFP(wp1872), lin-15(+)*]*; lin-15(n765)*. ZW1406: *zwEx262*[*Prab-3::adr-1::GFP(p1374), Pmyo-2::mStrawberry (wp1613)*]*; adr-1(zw96)*. ZW1407: *zwIs139*[*Pslo-1::slo-2(gf)(wp1311), Pmyo-2::YFP(wp214)*]*; adr-1(zw96)*. ZW1408: *zwIs139*[*Pslo-1::slo-2(gf)(wp1311), Pmyo-2::YFP(wp214)*]*; zwEx262*[*Prab-3::adr-1::GFP(p1374);Pmyo-2::mStrawberry (wp1613)*]*; adr-1(zw96)*. ZW1409: *scyl-1(zw99).* ZW1410: *slo-2(nf101); scyl-1(zw99).* ZW1415: *zwEx221*[*Prab-3::slo-2::GFP*]*; scyl-1(zw99)*. ZW1416: *zwEx247*[*Pslo-2::mStrawberry(wp1776), lin-15(+)*]*; zwEx263*[*Pscyl-1::GFP(wp1901+wp1902), lin-15(+)*]*; lin-15(n765).* ZW1417: *zwEx264*[*Prab-3::scyl-1(wp1912), Pmyo-2::mStrawberry (wp1613)*]; *scyl-1(zw99)*. ZW1418: *zwEx247*[*Pslo-2::mStrawberry(wp1776), lin-15(+)*]*; zwEx261*[*Padr-1::GFP(wp1872), lin-15(+)*]*; lin-15(n765)*. ZW1419: *zwEx265*[*Prab-3::GFP::scyl-1 3’UTR(wp1923), lin-15(+)*]*; lin-15(n765)*. ZW1420: *zwEx266*[*Prab-3::GFP::scyl-1 3’UTR(A-to-G)(wp1924), lin-15(+)*]*; lin-15(n765).* ZW1428: *slo-2(nf101); adr-1(zw96)*. ZW1505: *zwEx273*[*Prab-3::scyl-1::YFPc(wp1952), Prab-3::slo-2::YFPa(wp1783), lin-15(+)*]*; lin-15(n765)*. ZW1506: *zwEx274*[*Prab-3::scyl-1::YFPc(wp1952), Prab-3::slo-2N::YFPa(wp1784), lin-15(+)*]*; lin-15(n765)*. ZW1507: *zwEx275*[*Prab-3::scyl-1::YFPc(wp1952), Prab-3::slo-2C::YFPa(wp1785), lin-15(+)*]*; lin-15(n765)*.

### Mutant screening and mapping

An integrated transgenic strain expressing P*slo-1*::SLO-2(*gf*) and P*myo-2*::YFP (transgenic marker) in the wild-type genetic background was used for mutant screen. L4-stage *slo-2(gf)* worms were treated with the chemical mutagen ethyl methanesulfonate (50 mM) for 4 hours at room temperature. F2 progeny from the mutagenized worms were screened under stereomicroscope for animals that moved better than the original *slo-2(gf)* worms. 17 suppressors were isolated in the screen and were subjected to whole-genome sequencing. Analysis of the whole-genome sequencing data showed that 2 mutants have mutations in the *adr-1 gene* (www.wormbase.com). Identification of *adr-1* mutants was confirmed by the recovery of the sluggish phenotype when a wild-type cDNA of *adr-1* under the control of P*rab-3* was expressed in *slo-2(gf);adr-1(zw81)* double mutants.

### Generation of adr-1 and scyl-1 knockout mutants

The CRISPR/Cas9 approach (Dickinson et al., 2013) was used to create *adr-1* and *scyl-1* knockouts. The guide RNA sequences for *adr-1* and *scyl-1* are 5’-CCAGTTTTCGAAGCTTCGG and 5’-GAGGAGATTGGAAAATTGG, which were inserted into pDD162 (P*eft-3::Cas9* + Empty *sgRNA*; Addgene #47549), respectively. The resultant plasmids (*wp1645* for *adr-1* and *wp1887* for *scyl-1*) were injected into wild type worms, respectively, along with a repair primer (5’-GAGAAGTATTCACCAGTTTTCGAAGCTTAATGAGTTCCAAAAGATCCAGAGATTCCCGAA for *adr-1*, and 5’-TTGTAACAGCCGGAGGAGATTGGAAAATCTAGCTGGTGGACTTCATTTGGTCACTGGATT for *scyl-1*) and P*myo-2::mStrawberry* (*wp1613*) as the transgenic marker. The *adr-1* knockout worms were identified by PCR using primers 5’-TCACCAGTTTTCGAAGCTTAATGA (forward) and 5’-TCTTCTGCTGGCTCACATTCA (reverse). The *scyl-1* knockout worms were identified by PCR using primers 5’-CCGAAGTCCCAATTCCCAT (forward) and 5’-CCAAATGAAGTCCACCAGCTAG (reverse). The knockout worms were confirmed by Sanger sequencing.

### Analysis of expression pattern and subcellular localization

The expression pattern of *adr-1* was assessed by expressing GFP under the control of 1.8-kb *adr-1* promoter (P*adr-1::GFP*, *wp1872*). Primers for cloning P*adr-1* are 5’-TAAGGTACCAAGGACACGTTGCATATGAAT (forward) and 5’-TTTACCGGTTGGCTGACATATTGTGGGA (reverse). Subcellular localization of ADR-1 was determined by fusing GFP to its carboxyl terminus and expressing the fusion protein under the control of P*rab-3* (P*rab-3::adr-1::GFP*, *wp1374*). Primers for cloning *adr-1* cDNA are 5’-AAAGCGGCCGCATGGATCAAAATCCTAACTACAA (forward) and 5’-TTTACCGGTCCATCGAAAGCAGCAAGAGTGAAG (reverse). A plasmid (*wp1749*) harboring P*rab-3::his-58::mStrawberry* serves as a nucleus marker. The expression pattern of *scyl-1* was assessed by an *in vivo* recombination approach. Specifically, a 0.5 kb fragment immediately upstream of *scyl-1* initiation site was cloned and fused to GFP using the primers 5’-AATCTGCAGCATCGGCACGAGAAGTACA (forward) and 5’-TTAGGATCCCTAAAAGTGATCGAAATTTA (reverse). The resultant plasmid (P*scyl-1::GFP*, *wp1902*) was linearized and co-injected with a linearized (fosmid WRM068bA03), which contains 32 kb of *scyl-1* upstream sequence and part of its coding region, into the *lin-15(n765)* strain along with a *lin-15* rescue plasmid to serve as a transformation marker. To assay the effect of the identified adenosine site at the 3’UTR of *scyl-1* on gene expression, a 5.1 kb genomic DNA fragment covering part of the *scyl-1* last exon and subsequent sequence was cloned and fused in-frame to GFP using the primers 5’-AATGCTAGCATGCAGGCTAGAAATGAAGCTCG (forward) and 5’-TATGGGCCCGAAATCAGCATCTTTGACGAA (reverse). To mimic the A-to-I editing at the identified specific site, a second plasmid was made by mutating the specific adenosine to guanosine in the above plasmid. The two resultant plasmids were injected into *lin-15(n765)*, respectively, with a *lin-15* rescue plasmid as the transgenic marker. Images of transgenic worms were taken with a digital CMOS camera (Hamamatsu, C11440-22CU) mounted on a Nikon TE2000-U inverted microscope equipped with EGFP/FITC and mCherry/Texas Red filter sets (49002 and 49008, Chroma Technology Corporation, Rockingham, VT, USA).

### Behavioral assay

Locomotion velocity was determined using an automated locomotion tracking system as described previously (Wang and Wang, 2013). Briefly, a single adult hermaphrodite was transferred to an NGM plate without food. After allowing ∼30 sec for recovery from the transfer, snapshots of the worm were taken at 15 frames per second for 30 s using a IMAGINGSOURCE camera (DMK37BUX273) mounted on a stereomicroscope (LEICA M165FC). The worm was constantly kept in the center of the view field with a motorized microscope stage (OptiScanTM ES111, Prior Scientific, Inc., Rockland, MA, USA). Both the camera and the motorized stage were controlled by a custom program running in MATLAB (The MathWorks, Inc., Natick, MA).

### RNA-seq and data analysis

Total RNA was extracted from young adult-stage worms using TRIzol Reagent (Invitrogen) and treated with TURBO DNase (Ambion). RNA-seq was performed by Novogene Corp. Sacramento, CA Raw reads ware filtered using Trim Galore software (http://www.bioinformatics.babraham.ac.uk/projects/trim_galore/) to remove reads containing adapters or reads of low quality. The filtered reads were mapped to C. elegans genome (*ce*11) using TopHat2 (Kim et al., 2013). The gene expression level is estimated by counting the reads that map to exons.

### Bimolecular fluorescence complementation (BiFC) assay

BiFC assays were performed by coexpressing SLO-2 and SCYL-1 tagged with the amino and carboxyl terminal portions of YFP (YFPa and YFPc), respectively, in neurons under the control of *rab-3* promoter (P*rab-3*). To assay which portion of SLO-2 may interact with SCYL-1, the full-length, N-terminal, and C-terminal portion of SLO-2 were fused with YFPa, respectively. The resultant plasmids (*wp1783*, P*rab-3*::SLO-2::YFPa; *wp1784*, P*rab-3*::SLO-2N::YFPa, and *wp1785*, P*rab-3*::SLO-2C::YFPa) were coinjected with P*rab-3*::SCYL-1::YFPc (*wp1952*), respectively, into *lin-15(n765)* strain. A *lin-15* rescue plasmid was also coinjected to serve as a transformation marker. Epifluorescence of the transgenic worms was visualized and imaged as described above.

### C. elegans electrophysiology

Adult hermaphrodites were used in all electrophysiological experiments. Worms were immobilized and dissected as described previously (Liu et al., 2007). Borosilicate glass pipettes were used as electrodes for recording whole-cell currents. Pipette tip resistance for recording muscle cell currents was 3-5 MΩ whereas that for recording motor neuron currents was ∼20 MΩ. The dissected worm preparation was treated briefly with collagenase and perfused with the extracellular solution for 5 to 10-fold of bath volume. Classical whole-cell configuration was obtained by applying a negative pressure to the recording pipette. Current- and voltage-clamp experiments were performed with a Multiclamp 700B amplifier (Molecular Devices, Sunnyvale, CA, USA) and the Clampex software (version 10, Molecular Devices). Data were sampled at a rate of 10 kHz after filtering at 2 kHz. Spontaneous membrane potential changes were recorded using the current-clamp technique without current injection. Motor neuron whole-cell outward currents were recorded by applying a series of voltage steps (−60 to +70 mV at 10-mV intervals, 1200 ms pulse duration) from a holding potential of −60 mV. Spontaneous PSCs were recorded from body-wall muscle cells at a holding potential of −60 mV. Two bath solutions and three pipette solutions were used in electrophysiological experiments as specified in figure legends. Bath solution I contained (in mM) 140 NaCl, 5 KCl, 5 CaCl_2_, 5 MgCl_2_, 11 dextrose and 5 HEPES (pH 7.2). Bath solution II contained (in mM) 100 K^+^ gluconate, 50 KCl, 1 Mg^2+^ gluconate, 0.1 Ca^2+^ gluconate and 10 HEPES (pH 7.2). Pipette solution I contained (in mM) 120 KCl, 20 KOH, 5 Tris, 0.25 CaCl_2_, 4 MgCl_2_, 36 sucrose, 5 EGTA, and 4 Na_2_ATP (pH 7.2). Pipette solution II differed from pipette solution I in that 120 KCl was substituted by K^+^ gluconate. Pipette solution III contained (in mM) 150 K^+^ gluconate, 1 Mg^2+^ gluconate and 10 HEPES (pH 7.2).

### Xenopus oocytes expression and electrophysiology

A construct containing human *Slack* cDNA (pOX + *hSlo2.2*, a gift from Dr. Salkoff) was linearized with Pvu I. The mouse *Scyl1* cDNA was amplified from a construct (MR210762, Origene) and cloned into an existing vector downstream of the T3 promoter. The resultant plasmid (*wp*1982) was linearized with NgoM4. Capped cRNAs were synthesized using the mMessage mMachine Kit (Ambion). Approximately 50 nl cRNA of either *Slack* alone (0.5 ng/nl) or *Slack* (0.5 ng/nl) plus *Scyl1* (0.5 ng/nl) was injected into each oocyte using a Drummond Nanoject II injector (Drummond Scientific). Injected oocytes were incubated at 18°C in ND96 medium (in mM): 96 NaCl, 2 KCl, 1.8 CaCl_2_, 1 MgCl_2_, 5 HEPES (pH 7.5). 2 to 3 days after cRNA injection, single channel recordings were made in inside-out patches with a Multiclamp 700B amplifier (Molecular Devices, Sunnyvale, CA, USA) and the Clampex software (version 10, Molecular Devices). Data were sampled at 10 kHz after filtering at 2 kHz. Bath solution contained (in mM) 60 NaCl, 40 KCl, 50 K^+^ gluconate, 10 KOH, 5 EGTA, and 5 HEPES (pH 7.2). Pipette solution contained (in mM) 100 K^+^ gluconate, 60 Na^+^ gluconate, 2 MgCl_2_, and 5 HEPES (pH 7.2).

### Data Analyses for Electrophysiology

Amplitudes of whole-cell currents in response to voltage steps were determined from the mean current during the last 100 ms of the 1200-ms voltage pulses using the Clampfit software. The duration and charge transfer of PSC bursts were quantified with Clampfit software (version 10, Molecular Devices) as previously described (Liu et al., 2013). The frequency of PSC bursts was counted manually. For single channel analysis, the QuB software (https://qub.mandelics.com/) was used to fit open and closed times to exponentials, and to quantify the τ values and relative areas of the fitted components, which were automatically determined by the software. The first 30 sec recording of each experiment was used in such analyses. Statistical comparisons were made with Origin Pro 2019 (OriginLab Corporation, Northampton, MA) using either *ANOVA* or unpaired *t*-test as specified in figure legends. *p* < 0.05 is considered to be statistically significant. The sample size (*n*) equals the number of cells or membrane patches analyzed. All values are shown as mean ± SE and data graphing was done with Origin Pro 2019.

## Acknowledgements

This work was supported by National Institute of Health (R01GM113004 to B.C, and 2R01MH085927 and 1R01NS109388 to Z.-W.W.). We thank Dr. Laurence Salkoff for the human *Slack* construct. Some strains were provided by the CGC, which is funded by NIH Office of Research Infrastructure Programs (P40 OD010440).

## Figure legends

**Figure 4 supplement 1.**
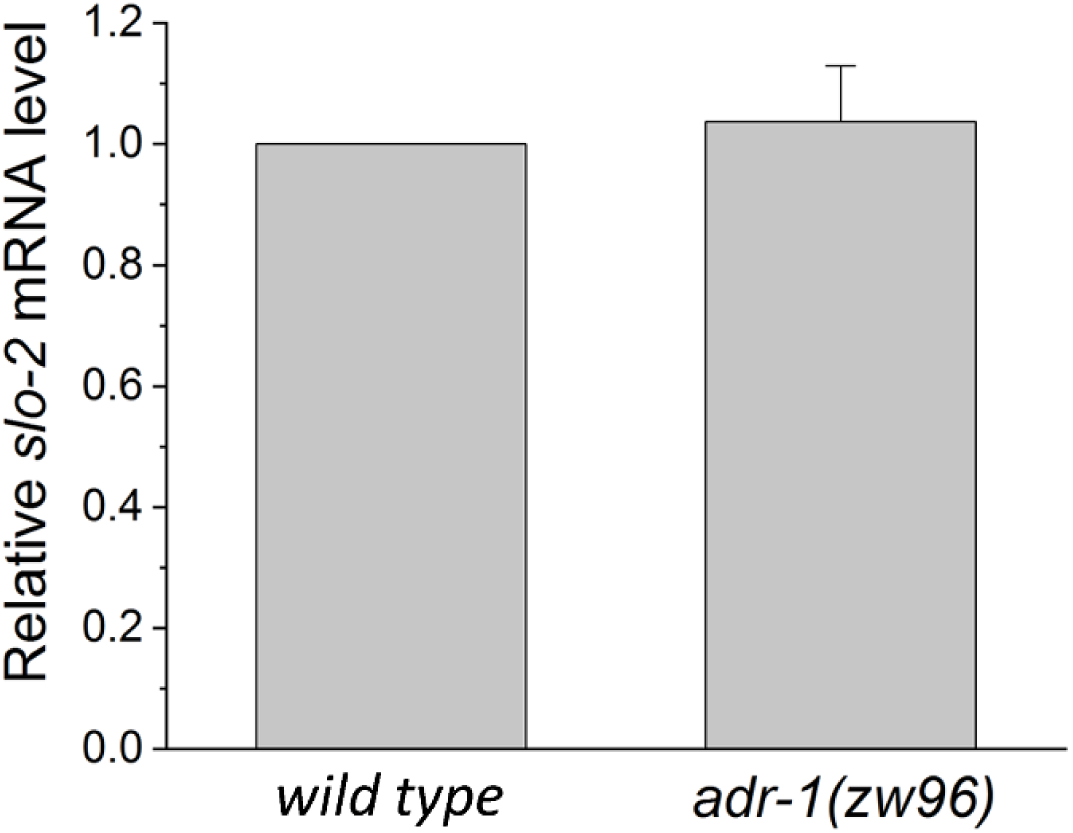
Comparison of *slo-2* transcript level between *wild type* and *adr-1* mutant. Shown are mean ± SE of three RNA-seq experiments. **Source data 1.** Raw data and numerical values for data plotted in Figure 4.

**Figure 5 supplement 1.**
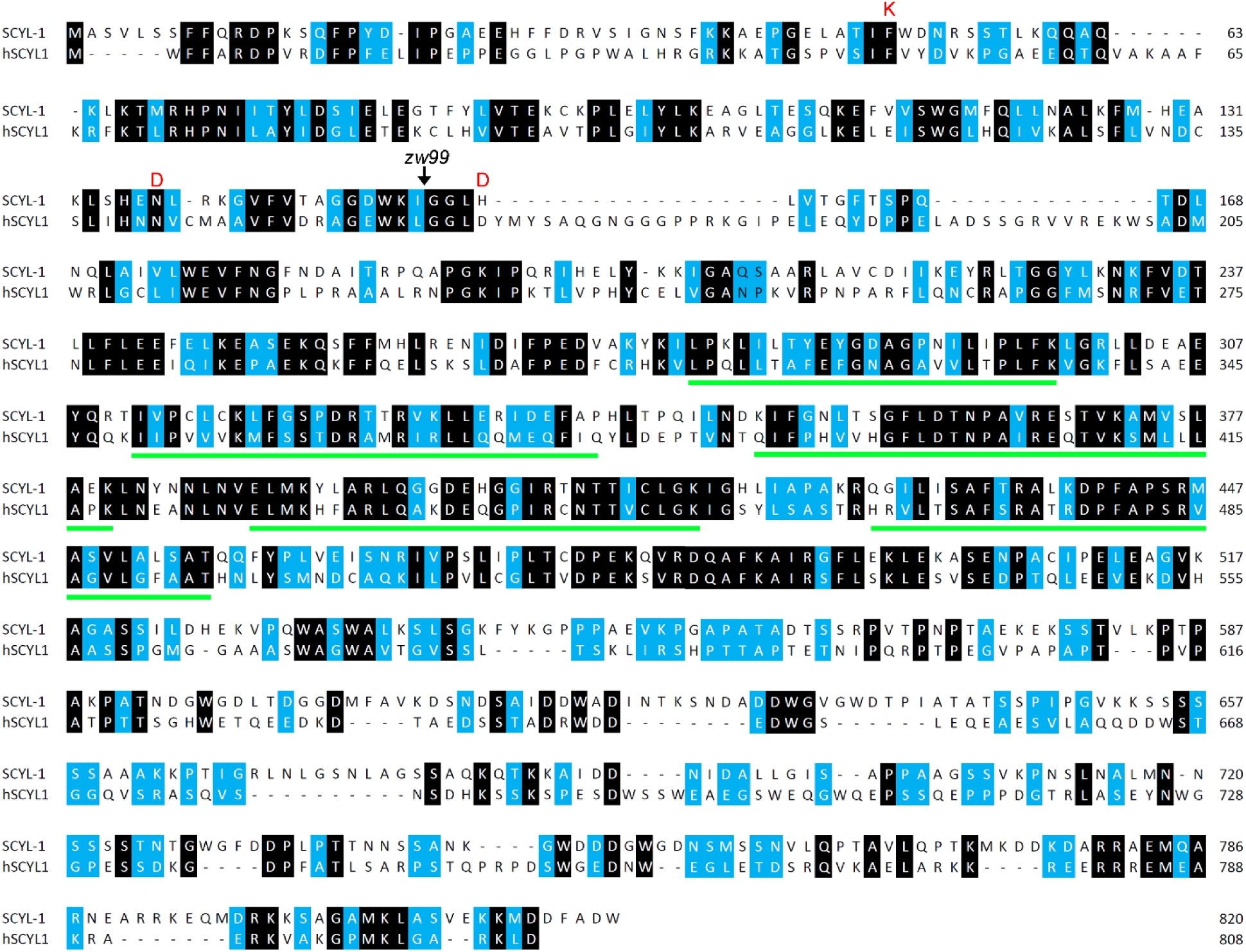
Alignment of amino acid sequences between *C. elegans* SCYL-1 (*W07G4.3*, www.wormbase.org) and human SCYL1 (hSCYL1, GenBank: NP_065731.3). Identical residues are highlighted in black, while similar ones (in size or polarity) in blue. The three residues that are essential for kinase activity in eukaryotic protein kinases are shown in red above the alignment at corresponding locations. Both proteins contain five HEAT repeats (marked by horizontal green lines) in the central portion. The *scyl-1* mutant allele *zw99* was made by introducing a stop codon after the residue I^152^ (indicated by an arrow) using the CRISPR/Cas9 approach. **Source data 1.** Raw data and numerical values for data plotted in Figure 5.

